# Selective Inhibitory Circuit Dysfunction after Chronic Frontal Lobe Contusion

**DOI:** 10.1101/2021.11.23.469731

**Authors:** Amber L. Nolan, Vikaas S. Sohal, Susanna Rosi

## Abstract

Traumatic brain injury (TBI) is a leading cause of neurologic disability; the most common deficits affect prefrontal cortex-dependent functions such as attention, working memory, social behavior, and mental flexibility. Despite this prevalence, little is known about the pathophysiology that develops in frontal cortical microcircuits after TBI. We investigated if alterations in subtype-specific inhibitory circuits are associated with cognitive inflexibility in a mouse model of frontal lobe contusion that recapitulates aberrant mental flexibility as measured by deficits in rule reversal learning. Using patch clamp recordings and optogenetic stimulation, we identified selective vulnerability in the non-fast spiking, somatostatin-expressing (SOM+) subtype of inhibitory neurons in layer V of the orbitofrontal cortex (OFC) two months after injury. These neurons exhibited reduced intrinsic excitability and a decrease in their synaptic output onto pyramidal neurons. By contrast, fast spiking, parvalbumin-expressing (PV+) interneurons did not show changes in intrinsic excitability or synaptic output. Impairments in SOM+ inhibitory circuit function were also associated with network hyperexcitability. These findings provide evidence for selective disruptions within specific inhibitory microcircuits that may guide the development of novel therapeutics for TBI.

## INTRODUCTION

Traumatic brain injury (TBI) is a leading cause of chronic neurologic disability (Engberg and Teasdale, 2004; Ponsford et al., 2008; Wilson et al., 2017). Damage often affects the frontal lobes, and thus the most frequent cognitive sequelae are deficits in prefrontal cortex (PFC)-dependent functions such as: attention, working memory, social behavior, mental flexibility, and task switching (Fujiwara et al., 2008; Spikman et al., 2012; Struchen et al., 2008; Stuss, 2011; Vilkki et al., 1994; Wallesch et al., 2001). Despite this prevalence, little is known about the chronic neuronal and network dysfunction associated with cognitive deficits after TBI, especially in frontal lobe circuits.

Cortical hyperexcitability has been identified in other regions of the neocortex adjacent to injury (Cantu et al., 2015; Carron et al., 2016; Ding et al., 2011). Loss of inhibitory function is postulated to contribute to these network changes due to a reduction in synaptic inhibition and decreases in expression of several immunohistochemical markers of interneurons, which have been reported particularly in severe TBI models (Buritica et al., 2009; Cantu *et al*., 2015; Carron *et al*., 2016; Nichols et al., 2018). However, the direct intrinsic and synaptic function of specific interneuron subtypes has not been systemically evaluated and compared.

Two major classes of GABAergic interneurons mediating inhibition of cortical output, particularly in deeper cortical layers, are parvalbumin-expressing (PV+) fast-spiking (FS) cells (45-60%), and somatostatin-expressing (SOM+) low threshold-spiking or regular spiking cells (20-50%) (Naka and Adesnik, 2016; Rudy et al., 2011). These neurons have distinct roles in the cortical microcircuit. PV+ neurons receive convergent thalamic input, synapse onto the perisomatic domains of pyramidal neurons, and have depressing synaptic inputs, supporting feedforward inhibition (Willems et al., 2018). This inhibition constrains the timing of temporal summation in pyramidal neurons (Pouille and Scanziani, 2001), affects stimulus tuning (Wang et al., 2004a; Zhu et al., 2015), increases the dynamic range of neuronal ensembles (Pouille et al., 2009), and mediates gamma oscillations during cognition (Cho et al., 2015). By contrast, SOM+ cells do not receive thalamic inputs, target dendrites instead of perisomatic regions, and have facilitating synapses (Naka and Adesnik, 2016). The combination of these properties promotes increasing SOM+ cell-mediated inhibition with increasing pyramidal neuron activity, producing feedback inhibition. While PV+ cells also provide an immediate feedback inhibition, SOM+ cells have been found to support a more delayed feedback pathway (Silberberg and Markram, 2007), that contributes to expansion of the input response range in sensory cortices (Murayama et al., 2009) as well as to synchronization of lower frequency theta and beta oscillations between circuits (Abbas et al., 2018; Chen et al., 2017). Changes in OFC inhibitory neurons, particular PV+ neurons, have been identified with reversal learning deficits in a sex-selective model of early life stress (Goodwill et al., 2018). How TBI alters these distinct inhibitory microcircuits in ways that may affect cortical circuit function and cognition has not been studied before.

We developed a mouse model of frontal lobe contusion that recapitulates cognitive inflexibility observed in humans after trauma (Chou et al., 2016). These mice exhibit deficits in reversal learning, a cognitive ability requiring orbitofrontal cortex (OFC) function (Bissonette et al., 2008). Interestingly, this type of contusion does not lesion the OFC, nor is the OFC in a directly pericontusional location. Thus, an immediately apparent mechanism for this deficit is unclear from the gross histology. The goal of this study was to understand if deficits in inhibition might be identified in the OFC at chronic time points after TBI when reversal learning impairments are observed. We utilized transgenic mouse lines and optogenetic techniques to systematically evaluate and compare the function of different subtypes of inhibitory interneurons in this model.

## RESULTS

### Non-fast spiking interneurons exhibit a selective reduction in intrinsic excitability at chronic time points after TBI

To investigate the neurophysiological changes that accompany the chronic reversal learning deficits which occur after frontal lobe contusion (Chou *et al*., 2016), we first assessed the intrinsic firing properties of layer V inhibitory neurons in the orbitofrontal cortex. Whole-cell patch clamp recordings were obtained from mice 2-3 months after undergoing a frontal lobe controlled cortical impact (CCI) or sham procedure. We visually identified interneurons using a transgenic mouse line that expresses a tdTomato fluorescent reporter in inhibitory neurons (*DlxI12b-Cre* x Ai14). A subset of neurons was filled with biocytin and processed for immunohistochemistry with somatostatin and parvalbumin antibodies after recording (Fig. 1A,K; n = 10 somatostatin positive, and n = 8 parvalbumin positive). The range of the action potential half-width and adaptation index in these immunohistochemically-verified somatostatin or parvalbumin-expressing neurons was then used to categorize recordings as fast spiking (FS, correlating with parvalbumin expression) or non-fast spiking (non-FS, correlating with somatostatin expression).

**Figure 1.**
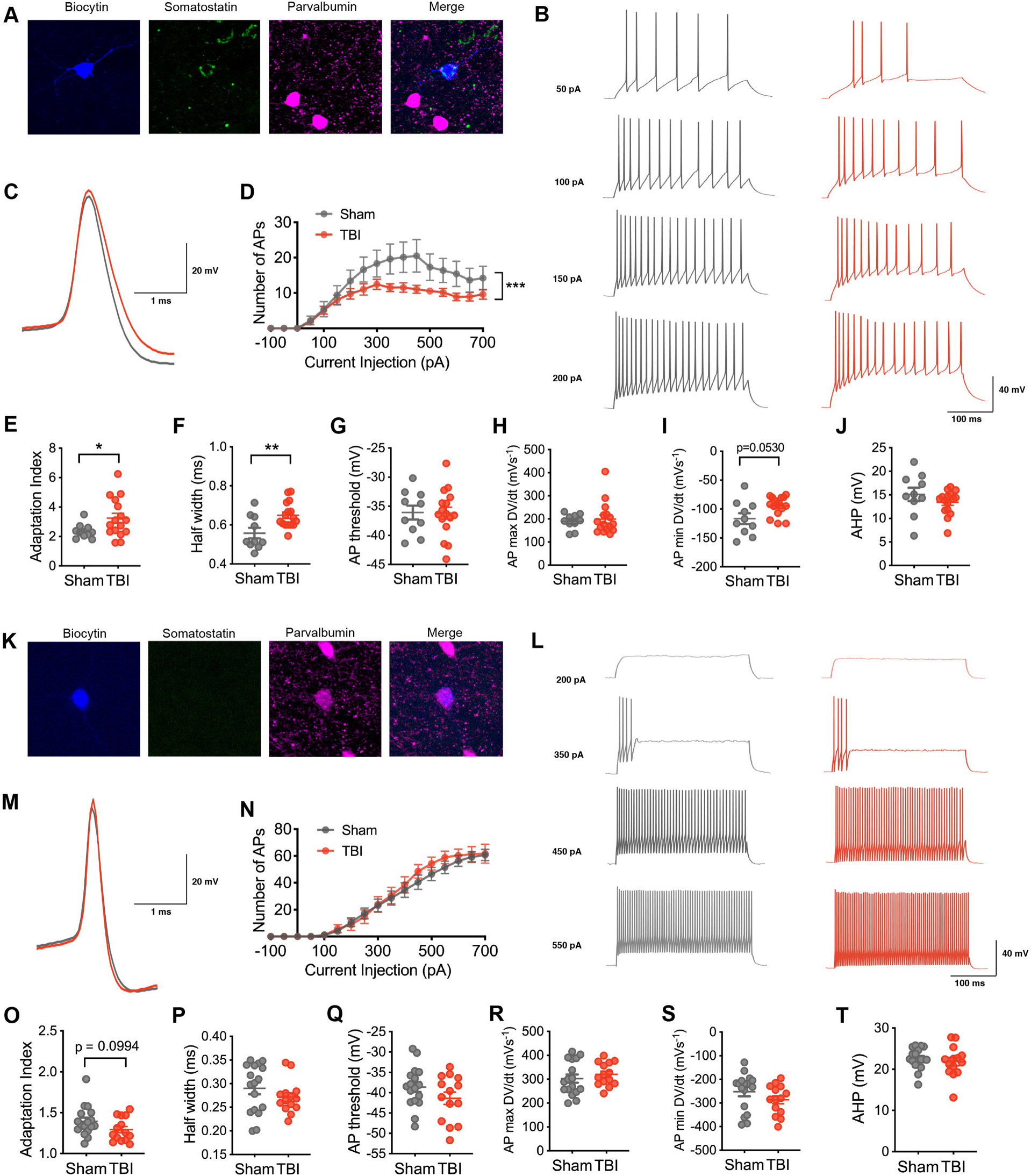
**Selective vulnerability of intrinsic excitability in non-fast spiking (SOM+) inhibitory neurons after chronic TBI.** A) A non-fast spiking interneuron in layer V of the orbitofrontal cortex that was filled with biocytin (blue) and later confirmed to express somatostatin (green) and not parvalbumin (pink). B) Representative current-clamp responses to depolarizing current steps in sham (grey) and TBI (red) mice. C) Average action potential shape. D) The number of action potentials plotted as a function of current injection (p = 0.0330 for TBI effect, and *** p = 0.0010 for interaction between current and TBI, repeated measures 2-way ANOVA). E) The adaptation index from current clamp responses measured at 400 pA above spiking threshold (* p = 0.0226, unpaired t-test with Welch’s correction). F-J) The action potential (AP) half-width, AP threshold, AP maximum rising slope, AP minimum falling slope, and afterhyperpolarization (AHP) calculated from current clamp responses 100 pA above spiking threshold (** p = 0.0053, unpaired t-test; p = 0.9529, unpaired t-test; p = 0.7366, Mann-Whitney test; p = 0.0530, unpaired t-test with Welch’s correction; p = 0.2323, unpaired t-test; respectively). K) A fast spiking interneuron in layer V of the orbitofrontal cortex that was filled with biocytin (blue) and later confirmed to express parvalbumin (pink) and not somatostatin (green). L) Representative current-clamp responses to depolarizing current steps in sham (grey) and TBI (red) mice. M) Average action potential shape. N) The number of action potentials plotted as a function of current injection (p = 0.6069 for TBI effect, repeated measures 2-way ANOVA). O) The adaptation index from current clamp responses measured at 400 pA above spiking threshold (p = 0.0994, unpaired t-test). P-T) The action potential (AP) half-width, AP threshold, AP maximum rising slope, AP minimum falling slope, and afterhyperpolarization (AHP) calculated from current clamp responses 100 pA above spiking threshold (p = 0.2574, unpaired t-test; p = 0.1614, unpaired t-test; p = 0.4253, unpaired t-test; p = 0.2010, unpaired t-test; p = 0.5087, unpaired t-test; respectively). Each neuron is represented with a symbol; solid lines indicate the mean ± SEM. (n=10 sham and 16 TBI non-fast spiking neurons from 7 (sham) and 8 (TBI) animals/group; n=17 sham and 14 TBI fast spiking neurons from 9 (sham) and 7 (TBI) animals/group).

Based on the response to a series of current injections, we evaluated intrinsic excitability in each inhibitory firing pattern subtype. Non-FS neurons from TBI animals exhibited a reduction in the number of action potentials seen with increasing current injection when compared to non-FS neurons from sham animals (Fig. 1B,D). This reduction in excitability was associated with a higher adaptation index (Fig. 1B,E) and a wider action potential half width (Fig. 1C,F). A non-significant trend was also noted for a reduced falling slope of the action potential (Fig. 1C, I). Other action potential metrics and passive membrane properties were not affected (Fig. 1G,H,J; Supplemental Table 1).

In contrast, intrinsic excitability was not altered in the FS group with TBI. A similar number of action potentials was found at varying amplitudes of current injection in both experimental groups (Fig. 1L,N). Other action potential and passive membrane properties were not significantly changed.

For comparison, we also evaluated the excitability of layer V pyramidal neurons. Subtle differences in action potential shape were found with TBI in comparison to neurons from sham animals (Supplemental Fig. 1C,H,J). However, these changes did not alter overall action potential production or intrinsic excitability (Supplemental Fig. 1B,D).

These findings demonstrate selective deficits in inhibitory neuronal function, specifically in the non-fast spiking/somatostatin interneuron subtype measured at chronic time points after frontal contusion injury.

### Optogenetic stimulation reveals subtype-specific deficits in inhibitory synaptic input

Next, we examined whether this intrinsic deficit was associated with a selective deficit in synaptic inhibition onto layer V pyramidal neurons at chronic time points after contusion. Towards this end, we optogenetically stimulated somatostatin-expressing (SOM+) or parvalbumin-expressing (PV+) inhibitory neurons with flashes of blue light (470 nm) across a high-power field and recorded optogenetically-evoked inhibitory post-synaptic currents (oIPSCs) in layer V pyramidal neurons (Fig. 2A, 2J). We expressed ChR2-EYFP in specific interneuron subtypes using transgenic mice (SOM-Cre x Ai32 and PV-Cre x Ai32 lines).

**Figure 2.**
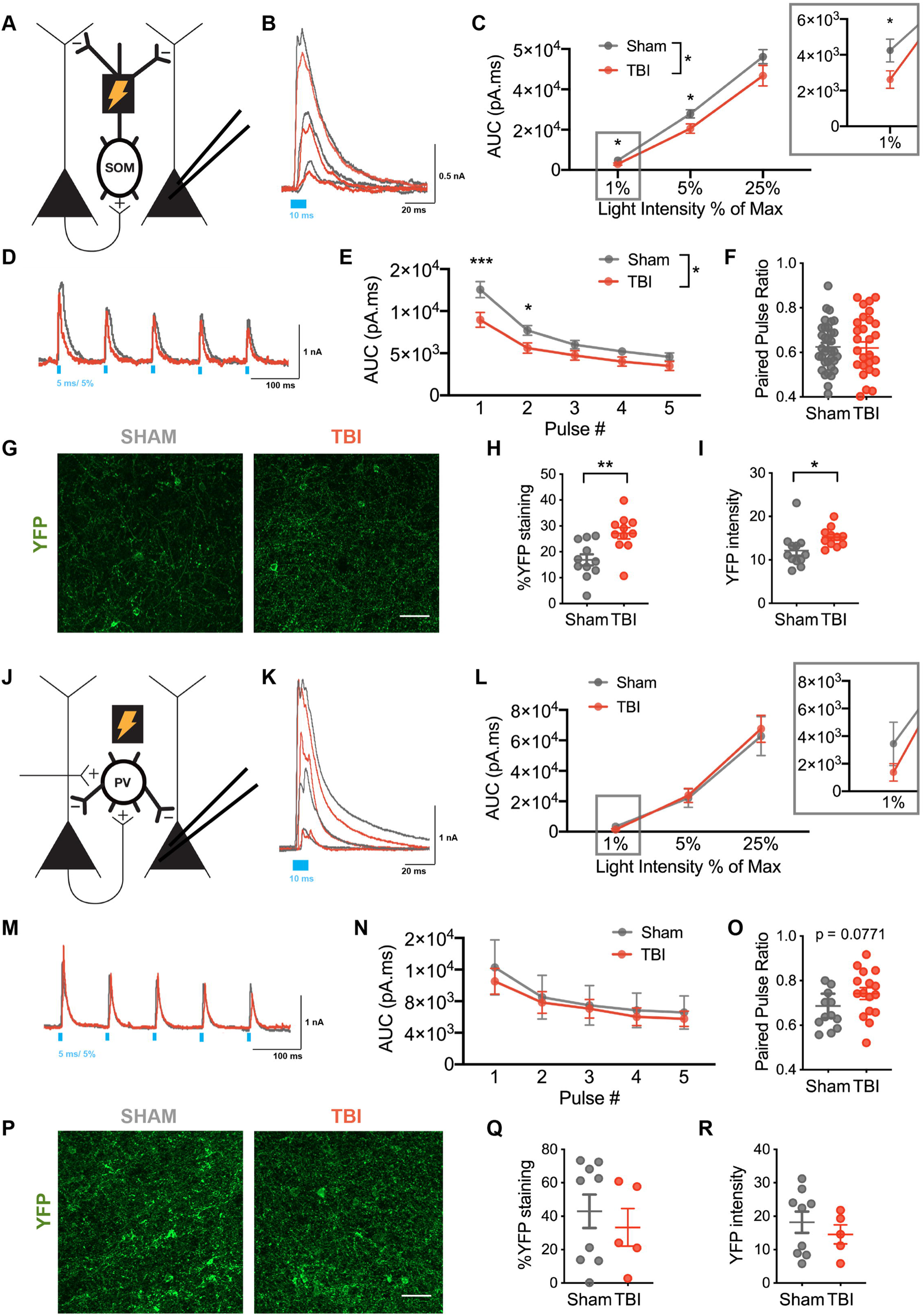
**Selective reduced SOM+-mediated inhibitory synaptic output on layer V pyramidal neurons after TBI.** A) Schematic of experimental design: voltage clamp recordings were obtained from layer V pyramidal neurons while activating ChR2-expressing SOM+ interneurons. B) Example optogenetically-evoked IPSCs (oIPSCs) recorded from pyramidal neurons in sham (grey) and TBI (red) conditions in response to 10 ms light pulses of increasing intensity (40 μW (1%), 225 μW (5%), and 1 mW/mm^2^ (25%)). C) Total charge (AUC) of oIPSC across light intensities (* p = 0.0347 for TBI effect, mixed effects model; * p < 0.05, post hoc tests controlling for the false discovery rate). D) Example oIPSCs in sham (grey) and TBI (red) conditions elicited with 10 Hz stimulation. E) Total charge (AUC) of oIPSCs elicited across pulse number (* p = 0.0281 for TBI effect, repeated measures two-way ANOVA; *** p < 0.0001, * p = 0.0189, post hoc tests controlling for the false discovery rate). F) The paired pulse ratio determined as the ratio of pulse 2/pulse 1 (p = 0.8864, unpaired t-test). G) Representative EYFP expression in layer V OFC in sham and TBI slices. H) Percentage area staining after binary thresholding (** p = 0.0032, unpaired t-test). I) Average EYFP intensity (* p = 0.0386, unpaired t-test). J) Schematic of experimental design: voltage clamp recordings were obtained from layer V pyramidal neurons while activating ChR2-expressing PV+ interneurons. K) Example optogenetically-evoked IPSCs (oIPSCs) recorded from pyramidal neurons in sham (grey) and TBI (red) conditions in response to 10 ms light pulses of increasing intensity (40 μW (1%), 225 μW (5%), and 1 mW/mm^2^ (25%)). L)Total charge (AUC) of oIPSC across light intensities (p = 0.8501 for TBI effect, repeated measures two-way ANOVA). M) Example oIPSCs in sham (grey) and TBI (red) conditions elicited with 10 Hz stimulation. N) Total charge (AUC) of oIPSC elicited across pulse number (p = 0.2649 for TBI effect, repeated measures two-way ANOVA). O) The paired pulse ratio determined as the ratio of pulse 2/pulse 1 (p = 0.0771; unpaired t-test). P) Representative EYFP expression in layer V OFC in sham and TBI slices. Q) Percentage area staining after binary thresholding (p = 0.5545 unpaired t-test). R) Average EYFP intensity (p = 0.4649, unpaired t-test). Circles represent the mean, solid lines indicate the SEM in C, E, L, N; Each neuron/slice is represented with a symbol and solid lines indicate the mean ± SEM in F, H, I, O, Q, R. (n=34 sham and 30 TBI neurons from 7 (sham) and 6 (TBI) animals/group for oIPSC data from SOM+ stimulation; n=12 sham and 11 TBI slices for immunohistochemistry for EYFP expression in SOM+ neurons from 4 (sham) and 4 (TBI) animals/group; n= 16 sham and 17 TBI neurons from 4 (sham) and 4 (TBI) animals/group for oIPSC data from PV+ stimulation; n= 9 sham and 5 TBI slices for immunohistochemistry for EYFP expression in PV+ neurons from 3 (sham) and 3 (TBI) animals/group).

During SOM-specific stimulation, oIPSCs were significantly reduced in neurons from TBI compared to sham animals as measured by the total charge elicited with light stimulation (Fig. 2B-C). Interestingly, post-hoc evaluation suggested that this reduction was most notable at lower light intensities, specifically 1% and 5% of the maximum light intensity. In addition to assessing the response to a unitary light pulse, we also determined the response to repetitive stimulation at various frequencies (10 and 40 Hz). Similar to the unitary pulse, repetitive light stimulation at the 5% light intensity demonstrated a lower response in neurons from TBI animals, most prominent with the first few pulses of light (Fig. 2D-E; Supplemental Fig. 2B-C). Also similar to the unitary pulse data, a TBI-associated reduction in response was less apparent at a higher light intensity (25%, Supplemental Fig. 3A-C). To assess whether these synaptic deficits were driven by changes in presynaptic release probability, we measured the frequency-dependent attenuation of oIPSCs with the paired-pulse ratio (PPR) at both 10 and 40 Hz stimulation. No change in the PPR was identified at either frequency to support TBI-induced chronic changes in the presynaptic release probability (Fig. 2F, Supplemental Fig. 2D, Supplemental Fig. 3D).

In contrast, during PV-specific stimulation, TBI did not alter the total charge of oIPSCs, whether with unitary (Fig. 2K-L) or repetitive light stimulation (Fig. 2M-N, Supplemental Fig. 2E-G, Supplemental Fig. 3E-G) at all tested light intensities. We did observe a non-significant trend towards an increase in the paired pulse ratio.

To rule out potential confounding reductions in ChR2-EYFP expression, we confirmed that EYFP fluorescence was not lower after TBI. Indeed, EYFP expression was not reduced in somatostatin neurons; surprisingly, it was increased in the TBI compared to sham group, both the percentage area of staining after binary thresholding, as well as the average intensity (Fig. 2G-I). Thus, the functional impairment measured in somatostatin-specific synaptic inhibition may be even greater than our data suggest given the TBI-associated compensation of increased EYFP-expression. EYFP expression in the PV+ population was not different for TBI vs. sham groups (Fig. 2P-R).

These data further support selective deficits in SOM+ neuronal function at chronic time points after TBI, specifically SOM-specific reduction in synaptic inhibition in the orbitofrontal cortex.

### TBI-induced reduction of SOM-mediated inhibition is also partially mediated by changes in quantal size and frequency

To further probe how specific changes in synaptic function might contribute to reduced synaptic output from SOM+ interneurons after TBI, we analyzed quantal IPSCs during optogenetically-evoked asynchronous release (qIPSCs, Fig. 3A). We replaced calcium with strontium in the extracellular solution to desynchronize the evoked release of neurotransmitter, allowing analysis of quantal events from stimulated synapses. Similar to miniature IPSCs, the frequency of qIPSCs is thought to reflect presynaptic properties, while the amplitude corresponds to postsynaptic changes.

**Figure 3.**
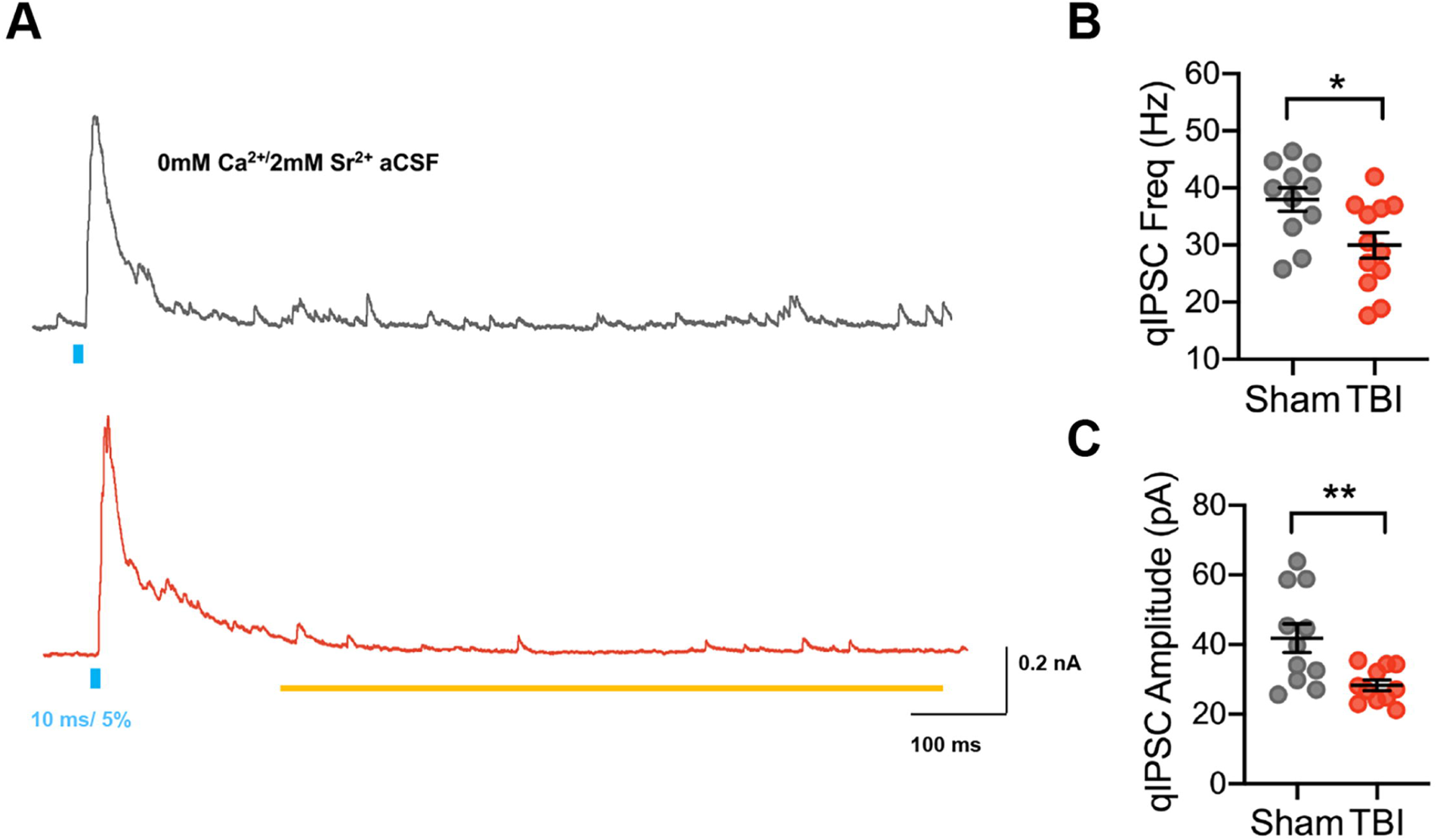
**TBI reduces both quantal size and frequency of SOM-mediated inhibition.** A) Example optogenetically-evoked qIPSCs elicited with strontium replacing calcium in the aCSF, recorded from pyramidal neurons in sham (grey) and TBI (red) conditions after 10 ms light stimulation of SOM+ neurons. The yellow line denotes window of analysis after stimulation. B) qIPSC frequency (* p = 0.0159; unpaired t-test). C) qIPSC amplitude (** p = 0.0086; Welch’s t-test). Each neuron is represented with a symbol and solid lines indicate the mean ± SEM. (n= 11 sham and 12 TBI neurons from 3 (sham) and 3 (TBI) animals/group).

SOM-specific optogenetic stimulation revealed that TBI altered both qIPSC frequency and amplitude at SOM+ inhibitory synapses when compared to sham recordings. Specifically, both the amplitude and frequency were reduced in TBI compared to sham cohorts (Fig. 3B,C). This suggests that changes in both pre- and post-synaptic factors may contribute to the reduced inhibitory output from SOM+ interneurons onto pyramidal neurons after TBI.

### Cortical network excitability is associated with a reduction of SOM-mediated inhibition after TBI

To assess if SOM-specific deficits in inhibition in the OFC correlate with a hyperexcitable cortical network, we measured the onset time to synchronized network input and the likelihood of large paroxysmal depolarization events in TBI and sham slices incubated in aCSF with 0 mM Mg^2+^ and 5 mM K^+^. In this solution, we observed the gradual development of polysynaptic synchronized events (> 2mV in amplitude) within current clamp recordings over time. In some cases, this progressed further to large, prolonged events (> 20 mV in amplitude, > 2 sec in duration) with overlaid high frequency spiking activity, termed paroxysmal depolarizations (PDS) (Fig. 4A).

**Figure 4.**
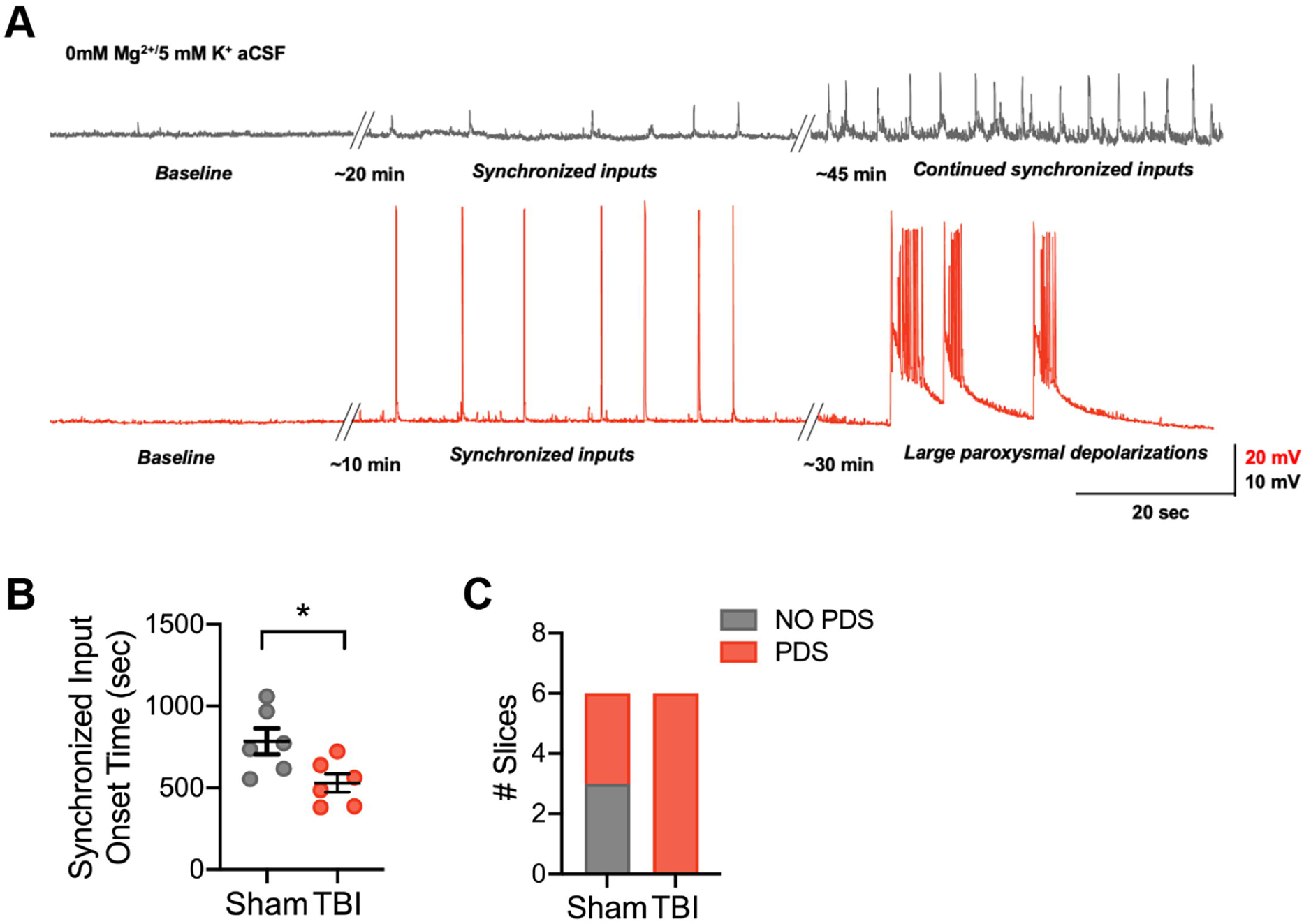
**Frontal lobe contusion induces network hyperexcitability in the orbitofrontal cortex.** A) Representative current clamp recordings from layer V pyramidal neurons of the OFC in a sham (grey) and TBI (red) animal showing synchronized input induced with a 0 mM Mg2+/5 mM K+ ACSF. Transition to large paroxysmal depolarizations with higher frequency components only occurred in the TBI animal in this example. B) The latency from 0 mM Mg2+/5 mM K+ ACSF application until the start of synchronized inputs (*, p = 0.0266, unpaired t-test). C) The number of slices that progressed to large paroxysmal depolarizations in TBI compared to sham slices (p = 0.0455, Chi Square test). Each neuron is represented with a symbol and solid lines indicate the mean ± SEM. (n= 6 sham and 6 TBI neurons from 2 (sham) and 3 (TBI) animals/group)

Slices from animals that underwent TBI 2-3 months prior had a significantly shortened latency to the onset of synchronized synaptic inputs compared to slices from sham animals (Fig. 4B). In addition, all slices from animals that received TBI exhibited PDS events within 45 minutes of incubation in this hyperexcitable solution, while only half of the sham slices showed this activity. These data support the hypothesis that TBI-induced SOM-specific disinhibition may alter the network balance of excitation and inhibition required for OFC function.

## DISCUSSION

Here we dissected inhibitory circuit function in the orbitofrontal cortex using a mouse model of frontal lobe contusion that exhibits deficits in reversal learning. Our findings demonstrate a selective vulnerability in the non-fast spiking, somatostatin-expressing subtype of inhibitory neurons at chronic time points after TBI. Specifically, these neurons exhibited a reduction in excitability and a decrease in their associated synaptic output onto pyramidal neurons. In contrast, fast spiking, parvalbumin-expressing neurons did not show changes in intrinsic excitability or synaptic output onto pyramidal neurons. Interestingly, this reduction in SOM-inhibitory network function was not due solely to changes in intrinsic excitability, as both the amplitude and frequency of quantal IPSCs originating from SOM interneurons were diminished after TBI. Finally, this selective vulnerability of SOM+ neuronal function identified in the OFC with chronic TBI was associated with a hyperexcitable network as measured by the onset time to synchronized polysynaptic input and an increased likelihood of large paroxysmal depolarizations. These findings support selective vulnerability of inhibitory populations in TBI and a possible target for therapeutic development.

Careful dissection and comparison of cortical inhibitory neuron subtypes and their circuit function has not been previously performed in the context of TBI, especially at chronic time points when persistent neurological deficits occur. Assessment of intrinsic firing patterns in particular has been limited. PV+/FS neuronal intrinsic function has been evaluated a few weeks after a severe frontoparietal pediatric TBI induced with a controlled cortical impact as well as at 24 hours after a mild central fluid percussion injury. Similar to our data, neither study found a change in intrinsic excitability (Nichols *et al*., 2018; Vascak et al., 2018), but the SOM+/non-FS population was not evaluated. Also, outside of the cortex, PV+ neurons in the dentate gyrus did not show changes in intrinsic membrane properties ~ 1 week after lateral fluid percussion injury (Folweiler et al., 2020).

A general reduction in synaptic inhibition has been observed in the cortex after TBI. Spontaneous inhibitory postsynaptic currents are reduced in frequency in layer V pyramidal neurons 2-4 weeks after a severe CCI centered over the sensorimotor cortex (Cantu *et al*., 2015). Interestingly, this reduction in sIPSC frequency has also been noted in FS interneurons in a severe pediatric CCI (Nichols *et al*., 2018) and acutely in mild TBI (Vascak *et al*., 2018). While the authors attribute this reduction to a loss of PV+ neurons or axonal processes, FS neurons also receive inhibition from somatostatin neurons (Cottam et al., 2013; Xu et al., 2013). These findings may thus be compatible with our finding showing selective reduction in subtype-specific inhibition onto layer V pyramidal neurons. Further research will be needed to interrogate if other neuronal types and lamina exhibit similar changes in SOM+ inhibitory control.

Our research supports that both changes in intrinsic and synaptic function underlie the reduction of SOM-mediated inhibition to pyramidal neurons. In addition to a change in intrinsic action potential production, both the frequency and amplitude of SOM-mediated qIPSCs were reduced after TBI. While we cannot completely exclude that a reduction in amplitude could affect the ability to identify events and artifactually reduce the detection frequency, these findings are potentially consistent with changes in both pre- and post-synaptic aspects of SOM-mediated inhibition. A reduction in inhibitory synapses has been found after TBI (Vascak et al., 2017), and a differential effect on GABA(A) receptor subunit types has been reported (Guerriero et al., 2015; Raible et al., 2012), which might correspond to changes in synapse number (pre-synaptic) and/or quantal size (post-synaptic). Why these changes appear limited to SOM+ synapses in our data is unclear. Axonal branches of SOM+ Martinotti neurons extend throughout the layers with extensive horizontal arborization, especially in layer I (Wang et al., 2004b). Perhaps this anatomy is more sensitive to compression from contusion after TBI, selectively reducing more distal dendritic input and the number of SOM+ synapses to pyramidal neurons. Interestingly, calretinin-expressing interneurons, of which a subset are SOM+ Martinotti neurons (Cauli et al., 2014; Nigro et al., 2018), are reported to selectively remodel neurite structure after injury (Brizuela et al., 2017), suggesting a capacity for maladaptive plasticity that selectively occurs in a subset of SOM+ neurons. Further dissection of specific pre- and post-synaptic mechanisms that mediate SOM-inhibition is needed to refine a more intricate understanding of inhibitory networks after TBI.

Previous research has often focused on a loss of inhibitory neurons after TBI as measured with immunohistochemistry (Buritica *et al*., 2009; Cantu *et al*., 2015; Nichols *et al*., 2018). Interestingly, comparison of immunohistochemistry with transgenic mouse lines, suggests that some of this reduction is secondary to changes in the expression of markers rather than an actual loss of neurons (Nichols *et al*., 2018). In our model, at chronic time points, we did not document a loss of PV+ or SOM+ neurons or processes as measured by expression of YFP in Cre-dependent transgenic mouse lines. Interestingly, while PV+ YFP expression is not affected, there was a paradoxical increase in SOM+ neuronal YFP expression. This finding might suggest a compensatory mechanism such as increased axonal sprouting in SOM+ neurons that is however, ineffective in fully correcting the chronic functional deficits observed here.

Maladaptive circuit changes within the OFC have not been widely investigated in animal models with deficits in reversal learning. Decreased expression and density of PV+ interneurons were identified in a model of early life stress that selectively impairs rule reversal learning in females. The authors demonstrated that optogenetically silencing PV+ neurons in the OFC also recapitulated these deficits in control mice (Goodwill *et al*., 2018). Another group using a mouse model of obsessive-compulsive disorder with deletion of the *Sapap3* gene reported reduced activity in GABAergic neurons of the OFC associated with deficits in reversal learning. Interestingly, optogenetically activating the GABAerigic neurons with 40 Hz stimulation, in the appropriate behavioral context (in which their activity was most reduced), rescued the behavioral deficits (Yang et al., 2021). Thus, inhibition is important for circuit function in reversal learning; however, exactly how SOM+ and PV+ neurons might interact to contribute to this behavior needs to be explored further.

More generally, impairment in SOM+ function is associated with pathological network activity and behavioral deficits. Mice lacking DLX1, a transcription factor important for the development of interneuron subsets, show a reduction of SOM+ but not PV+ cells, develop delayed-onset epilepsy and have deficits in behavioral inhibition and associative fear learning (Cobos et al., 2005; Mao et al., 2009). In epileptic circuits, loss of inhibitory input specifically to the dendritic compartment rather than perisomatic has been noted (Cossart et al., 2001), suggesting a selective dysfunction of SOM+ inhibition. Specific manipulation of SOM+ neurons through selective mutation of MeCP2 leads to repetitive stereotyped behaviors as well as seizures indicating a specific role for SOM interneurons in Rett syndrome (Ito-Ishida et al., 2015). Finally, in models of a pediatric genetic epilepsy with behavioral co-morbidities (Dravet’s syndrome, a sodium channel mutation), SOM+ (and PV+) neurons have a reduction in excitability that contributes to impairment in frequency-dependent di-synaptic inhibition (Tai et al., 2014) as well as autism-related behavioral phenotypes and deficits in spatial learning (Griffin et al., 2018).

It should be emphasized that somatostatin neurons are not a homogeneous population. In fact, these neurons represent a diverse group with heterogenous morphology and function. One “classic” type of SOM+ neuron, which is thought to comprise the majority of deeper layer SOM+ cells, is the Martinotti cell (MC). This specific subtype is characterized by translaminar ascending axon collaterals, especially in layer I, and low-threshold or regular spiking firing behavior (Riedemann, 2019). Other SOM+ neurons are often referred to as non-MCs and include morphologically defined basket cells, double-bouquet cells and long-range inhibitory projection cells amongst others (Riedemann, 2019). Basket cells may behave more similarly to PV+ neurons, exhibiting innervation around pyramidal cell bodies and quasi-fast spiking physiology (Naka et al., 2019; Riedemann, 2019). The non-MCs also have much higher spike thresholds compared to MC neurons (Naka *et al*., 2019; Nigro *et al*., 2018). Interestingly, our data suggests more deficits in SOM-specific input to pyramidal neurons at lower light intensities which should correspond to neurons with lower thresholds for action potential production. This might be indicative of specific dysfunction within MC cell populations after TBI.

In summary, we have established that SOM+ neurons are particularly vulnerable to TBI, exhibiting changes in intrinsic firing and synaptic output to pyramidal neurons, while in another major class of PV+ inhibitory neurons, neuronal firing and synaptic output is relatively unaffected. Why these SOM+ interneurons, and exactly which subpopulation within this class, are particularly vulnerable is a question of great interest. The intrinsic ability for plasticity (Brizuela *et al*., 2017), unique axonal structure (Wang *et al*., 2004b), and/or interaction with neuroinflammatory mediators such as microglia and astrocytes (Um, 2017) needs to be fully investigated to inform potential subtype-targeted therapies for cognitive dysfunction and other sequelae of TBI.

## Supporting information

Supplemental Key Resource Table

## ACKNOWLEDGEMENTS

A.L.N. was supported by NIH/NINDS (K08NS114170) and the UCSF Clinical and National Center for Advanced Translational Sciences at NIH (UCSF-CTSI Grant Number TL1 TR001871). S.R. was supported by NIH/National Institute on Aging Grant R01AG056770).

## AUTHOR CONTRIBUTIONS

A.L.N. performed the experiments and analysis. A.L.N., V.S.S., and S.R. designed the experiments and wrote the paper.

## DECLARATION OF INTERESTS

The authors declare no competing interests.

## METHODS

### RESOURCE AVAILABILITY

#### Lead contact

Further information and requests for resources should be directed to and will be fulfilled by the lead contact, Amber L. Nolan.

#### Materials availability

This study did not generate new unique reagents.

#### Data and code availability

All data reported in this paper will be shared by the lead contact upon request.

This paper does not report original code.

Any additional information required to reanalyze the data reported in this paper is available from the lead contact upon request.

### EXPERIMENTAL MODEL AND SUBJECT DETAILS

#### Mice

All experiments were conducted in accordance with National Institutes of Health (NIH) Guide for the Care and Use of Laboratory Animals and approved by the Institutional Animal Care and Use Committee of the University of California, San Francisco (AN184326-02B).

Male and female mice were bred in house using the following strains: B6.129P2-Pvalb^tm1(cre)Arbr^/J (PV-Cre, The Jackson Laboratory), Sst^tm2.1(cre)Zjh/^J (SOM-Cre, The Jackson Laboratory), Tg(I12b-cre)1Jlr (DlxI12b-Cre, from Dr. John Rubenstein), B6.Cg-Gt(ROSA)26Sor^tm14(CAG-tdTomato)Hze^/J (Ai14, The Jackson Laboratory), B6.Cg-Gt(ROSA)26Sor^tm32(CAG-COP4*H134R/EYFP)Hze^/J (Ai32, The Jackson Laboratory). Mice were group housed in environmentally controlled conditions with a 12:12 h light: dark cycle at 21 ± 1 °C; ~50% humidity and provided food and water ad libitum. Mice were 12-13 weeks of age at the time of surgery. Both males and females were used and results from both sexes were pooled. We did not observe any influence or association of sex on the experimental outcomes.

### METHOD DETAILS

#### Surgical Procedures

Mice were randomly assigned to a TBI or sham surgery group. They were anesthetized and maintained under anesthesia at 2-2.5% isoflurane. TBI, specifically frontal lobe contusion, was achieved using a controlled cortical impact (CCI) surgery, performed as previously described (Chou *et al*., 2016). Briefly, mice were secured in a stereotaxic frame. A midline incision exposed the skull followed by a ~2.5 mm diameter craniectomy at +2.34 mm anteroposterior and +1.62 mm mediolateral with respect to bregma. After the craniectomy, contusion was induced using a 2 mm convex tip attached to an electromagnetic impactor (Leica). The contusion depth was set to 1.25 mm from dura with a velocity of 4.0 m/s sustained for 300 ms. Following impact, the scalp was sutured. These injury parameters were chosen to induce contusion over the right frontal lobe, but not result in a large cavity. Sham animals were subjected to identical parameters excluding the craniectomy and impact.

#### Slice Preparation

Sagittal brain slices (250 µm) including the orbitofrontal cortex (between ~ +0.7 mm to ~ +1.7 mm from bregma) were prepared from mice that underwent TBI or a sham procedure, ~2-3 months prior. Mice were anesthetized with Euthasol (0.1ml/25g, Virbac, Fort Worth, TX, NDC-051311-050-01), and transcardially perfused with an ice-cold sucrose solution containing (in mM): 210 sucrose, 1.25 NaH_2_PO_4_, 25 NaHCO_3_, 2.5 KCl, 0.5 CaCl_2_, 7 MgCl_2_, 7 dextrose, 1.3 ascorbic acid, 3 sodium pyruvate (bubbled with 95% O2−5% CO2, pH 7.4). Mice were then decapitated, and the brain was isolated in the same sucrose solution, and cut on a slicing vibratome (Leica, VT1200S, Leica Microsystems, Wetzlar, Germany). Slices were incubated in a holding solution (composed of (in mM) 125 NaCl, 2.5 KCl, 1.25 NaH_2_PO_4_, 25 NaHCO_3_, 2 CaCl_2_, 2 MgCl_2_, 10 dextrose, 1.3 ascorbic acid, 3 sodium pyruvate, bubbled with 95% O2−5% CO2, pH 7.4) at 36°C for 30 minutes and then at room temperature for at least 30 minutes until recording.

#### Intracellular Recording

Whole cell recordings were obtained from these slices in a submersion chamber with a heated (32–34°C) artificial cerebrospinal fluid (aCSF) containing (in mM): 125 NaCl, 3 KCl, 1.25 NaH_2_PO_4_, 25 NaHCO_3_, 2 CaCl_2_, 1 MgCl_2_, 10 dextrose (bubbled with 95% O2/5% CO2, pH 7.4). Patch pipettes (3–6MΩ) were manufactured from filamented borosilicate glass capillaries (Sutter Instruments, Novato, CA, BF100-58-10) and filled with an intracellular solution containing (in mM): 135 KGluconate, 5 KCl, 10 HEPES, 4 NaCl, 4 MgATP, 0.3 Na_3_GTP, 7 2K-phosphcreatine, and 1-2% biocytin. For recording of qIPSCs, a cesium-based intracellular solution was instead used containing (in mM): 135 CsMeS, 5 CsCl, 10 HEPES, 4 NaCl, 4 MgATP, 0.3 Na_3_GTP, 7 2K-phosphcreatine, 2 Qx314Br, and 1-2% biocytin. Neurons were visualized using infrared microscopy with a 40x water-immersion objective (Olympus, Burlingame, CA). Pyramidal neurons and tdTomato expressing inhibitory neurons located in layer V were targeted for patching. Recordings were made using a Multiclamp 700B (Molecular Devices, San Jose, CA) amplifier, which was connected to the computer with a Digidata 1440A ADC (Molecular Devices, San Jose, CA), and recorded at a sampling rate of 20 kHz with pClamp software (Molecular Devices, San Jose, CA). We did not correct for the junction potential, but pipette capacitance was appropriately compensated before each recording.

The passive membrane and active action potential spiking characteristics were assessed by injection of a series of hyperpolarizing and depolarizing current steps with a duration of 250 ms from −250 pA to 700 nA (in increments of 50 pA). The resting membrane potential was the measured voltage of the cell after obtaining whole cell configuration without current injection. A holding current was then applied to maintain the neuron at ~ −67 +/− 3 mV before/after current injections. The input resistance was determined from the steady-state voltage reached during the −50 pA current injection. The membrane time constant was the time required to reach 63% of the maximum change in voltage for the −50 pA current injection.

Action potential (AP) properties including the half width, threshold, amplitude, rising slope, falling slope, and spike afterhyperpolarization (AHP) were calculated based on the response to a current pulse that was 100 pA above the minimal level that elicited spiking, or if this injection had fewer than 3 action potentials, the first current injection with >3 action potentials. The AP threshold was defined as the voltage at which the third derivative of V (d^3^V/dt) was maximal just prior to the action potential peak. The AP amplitude was calculated by measuring the voltage difference between the peak voltage of the action potential and the spike threshold. The half-width was determined as the duration of the action potential at half the amplitude. The spike AHP was the voltage difference between the AP threshold and the minimum voltage before the next AP. The rising and falling slopes of the AP were the maximum of the first derivative of V between the threshold to the peak amplitude, and the peak amplitude to the minimum voltage before the next AP, respectively.

Action potential timing was detected by recording the time at which the positive slope of the membrane potential crossed −20 mV. From the action potential times, the instantaneous frequency for each action potential was determined (1/inter spike interval). The adaptation index of each cell was the ratio of the first over the last instantaneous firing frequency in response to a current pulse that was 400 pA above the minimal level that elicited spiking. The AP rate as a function of current injection was examined by plotting the first instantaneous AP frequency versus current injection amplitude. The rheobase was then calculated as the intercept of the best linear fit of the first four positive values of this plot.

As noted above, we identified interneurons based on the expression of tdTomato driven by a Cre-dependent fluorescent reporter mouse line (Ai14) crossed to a DlxI12b-Cre mouse (Potter et al., 2009). This strategy labels a majority of interneuron subtypes. Recorded interneurons were therefore subdivided into fast-spiking (FS) or non-FS groups based on the range of electrophysiological properties identified in a subset of these neurons that were successfully filled with biocytin and processed for immunohistochemistry with somatostatin and parvalbumin antibodies after recording. The criteria for the FS group was an adaptation index of <1.9 and a half width of <0.35 msec based on the 8 histologically verified parvalbumin-expressing neurons. The criteria for the non-FS group was an adaptation index of >1.6 and a half width of 0.45-0.8 msec based on the 10 histologically verified somatostatin-expressing neurons.

#### ChR2 Stimulation

To measure optogenetically-evoked SOM+- or PV+-mediated inhibitory post synaptic currents (oIPSC), SOM-Cre and PV-Cre mice were crossed with a channelrhodopsin-2 (ChR2)/EYFP reporter line (Ai32). Voltage clamp recordings were attained from layer V pyramidal neurons in the OFC while the cell was held at 0 mV in voltage clamp. 10 msec flashes of blue light, generated by a Lambda DG-4 high-speed optical switch with a 300 W Xenon lamp (Sutter Instruments) and an excitation filter set centered around 470 nm, were delivered to the slice through a 40x objective (Olympus) to activate ChR2 at a range of light intensities from 40 uW/mm^2^ to 1 mW/mm^2^ (1-25% of the maximum intensity generated). At each light intensity, the light pulse was repeated five times with a 4 sec interval in between each stimulus. The five oIPSCs were averaged and the total charge/ area under the curve (AUC), for 100 msec after the light pulse, was measured for each light intensity. Using a similar protocol, we also delivered trains of 5 ms light pulses at both 10 and 40 Hz stimulation frequencies. The AUC between light pulses was determined for each pulse in the train of pulses. The paired pulse ratio was calculated as the ratio of the response to the second pulse / first pulse.

To examine light-evoked quantal IPSCs (qIPSCs) specifically from SOM+ neurons, the calcium in the aCSF was replaced with an equal amount of strontium (2 mM SrCl_2_ instead of 2 mM CaCl_2_). As described above, voltage clamp recordings were obtained from layer V pyramidal neurons in SOM-CrexAi32 mice while activating ChR2 with a 10 msec light pulse delivered at the 5% light intensity (225 μW/mm^2^). The light pulse was repeated 30 times with a 10 sec interval in between each stimulus. Analysis of quantal events was performed over an 800 msec window beginning 200 msec after the stimulus onset, a time window designed to eliminate synchronous synaptic responses. qIPSC frequency and amplitude were analyzed using a template matching algorithm in ClampFit 10.7 (Molecular Devices, San Jose, CA). The template was created using recordings from multiple pyramidal cells and included several hundred qIPSC events. The first 200 qIPSCs or all the events measured in the 30 sweeps from each cell were included for analysis. The mean frequency and amplitude for these events was calculated for each cell to compare between groups.

#### Network Excitability

To evaluate network excitability in TBI and sham cortical networks, slices were incubated in aCSF without Mg^2+^ (0mM MgCl_2_) and with increased K^+^ (5 mM KCl). In this solution, synchronized polysynaptic events occur with time, and the time to onset of the second clearly polysynaptic burst with an amplitude > 2mV was compared while recording in current clamp from layer V neurons while holding the membrane potential to ~−65 to −70 mV with holding current. Activity was recorded for 45 minutes, and the occurrence of large paroxysmal depolarization events was also noted. These events were defined as input > 20 mV and lasting > 2 sec.

#### Immunohistochemistry

Brain slices obtained from electrophysiological recordings were drop-fixed in 4% paraformaldehyde in a phosphate-buffered solution (4–48 h), then rinsed with phosphate-buffered saline (PBS) with sodium azide (0.1 M PBS +0.05% NaN_3_) and stored at 4°C for 0–5 days until processing. Sections were then washed in PBS (3L L min), incubated in a blocking solution (10% donkey serum diluted in 0.5% Triton X-100 in 0.1M PBS) for 1 h, and then incubated with primary antibodies, somatostatin (1:1000, Abcam, ab64053) and parvalbumin (1:8000, Millipore, MAB1572) with 0.25% Triton X-100, and 5% donkey serum in 0.1 M PBS, at room temperature overnight, along with streptavidin-conjugated Pacific Blue (1:500, ThermoFisher Scientific, S11222). The following day, sections were then rinsed with PBS (3L L min) and incubated in secondary antibodies (1:500, Alexa 488 donkey anti-rabbit, Invitrogen, A21206, and 1:500, Alexa 647 donkey anti-mouse, Abcam, ab181292) at room temperature overnight. Finally, sections were rinsed and mounted in an aqueous medium. Images were obtained using a Zeiss Imager Z1 Apotome microscope controlled by ZEN software (Zeiss 2012).

For analysis of EYFP expression, slices were fixed as described above after recording optogenetically-evoked IPSCs from both SOM-CrexAi32 and PV-CrexAi32 mice. These slices were incubated with a primary antibody to enhance expression of EYFP (anti-GFP, 1:500, Exbio, 11-476), followed by secondary antibody (1:500, Alexa 488 donkey anti-rabbit, Invitrogen, A21206) as described above. The OFC was imaged with a 20x objective, centered over layer V obtaining a Z-stack of 15 μm with a 0.5-μm slice interval using the same exposure time across slices. Image stacks were flattened and analyzed using ImageJ (National Institutes of Health, Bethesda, MD). Both the average intensity as well as the percentage area of staining after binary thresholding using the same threshold across images were calculated.

### QUANTIFICATION AND STATISTICAL ANALYSIS

All data were evaluated with GraphPad Prism 8 statistical software. Statistical significance between groups for most variables was determined using an unpaired t-test with or without Welch’s correction. For nonparametric data, a Mann-Whitney test was assessed to determine significance. The firing responses to increasing current injections and the oIPSC AUC with increasing light intensity or pulse number were analyzed as a repeated measures two-way ANOVA or mixed effects model. Post-hoc multiple comparisons were assessed controlling for the false discovery rate using the method of Benjamini and Hochberg. A Chi Square test evaluated the probability of PDS occurrence. p values < 0.05 were considered significant. All of the statistical details of experiments can be found in the figure legends including the statistical tests used, exact value of n, and what n represents.

## Notes

### Competing Interest Statement

The authors have declared no competing interest.

## REFERENCES

Abbas, A.I., Sundiang, M.J.M., Henoch, B., Morton, M.P., Bolkan, S.S., Park, A.J., Harris, A.Z., Kellendonk, C., and Gordon, J.A. (2018). Somatostatin Interneurons Facilitate Hippocampal-Prefrontal Synchrony and Prefrontal Spatial Encoding. Neuron 100, 926–939 e923. 10.1016/j.neuron.2018.09.029.

Bissonette, G.B., Martins, G.J., Franz, T.M., Harper, E.S., Schoenbaum, G., and Powell, E.M. (2008). Double dissociation of the effects of medial and orbital prefrontal cortical lesions on attentional and affective shifts in mice. J Neurosci 28, 11124–11130. 10.1523/JNEUROSCI.2820-08.2008.

Brizuela, M., Blizzard, C.A., Chuckowree, J.A., Pitman, K.A., Young, K.M., and Dickson, T. (2017). Mild Traumatic Brain Injury Leads to Decreased Inhibition and a Differential Response of Calretinin Positive Interneurons in the Injured Cortex. J Neurotrauma 34, 2504–2517. 10.1089/neu.2017.4977.

Buritica, E., Villamil, L., Guzman, F., Escobar, M.I., Garcia-Cairasco, N., and Pimienta, H.J. (2009). Changes in calcium-binding protein expression in human cortical contusion tissue. J Neurotrauma 26, 2145–2155. 10.1089/neu.2009.0894.

Cantu, D., Walker, K., Andresen, L., Taylor-Weiner, A., Hampton, D., Tesco, G., and Dulla, C.G. (2015). Traumatic Brain Injury Increases Cortical Glutamate Network Activity by Compromising GABAergic Control. Cereb Cortex 25, 2306–2320. 10.1093/cercor/bhu041.

Carron, S.F., Alwis, D.S., and Rajan, R. (2016). Traumatic Brain Injury and Neuronal Functionality Changes in Sensory Cortex. Front Syst Neurosci 10, 47. 10.3389/fnsys.2016.00047.

Cauli, B., Zhou, X., Tricoire, L., Toussay, X., and Staiger, J.F. (2014). Revisiting enigmatic cortical calretinin-expressing interneurons. Front Neuroanat 8, 52. 10.3389/fnana.2014.00052.

Chen, G., Zhang, Y., Li, X., Zhao, X., Ye, Q., Lin, Y., Tao, H.W., Rasch, M.J., and Zhang, X. (2017). Distinct Inhibitory Circuits Orchestrate Cortical beta and gamma Band Oscillations. Neuron 96, 1403–1418 e1406. 10.1016/j.neuron.2017.11.033.

Cho, K.K., Hoch, R., Lee, A.T., Patel, T., Rubenstein, J.L., and Sohal, V.S. (2015). Gamma rhythms link prefrontal interneuron dysfunction with cognitive inflexibility in Dlx5/6(+/−) mice. Neuron 85, 1332–1343. 10.1016/j.neuron.2015.02.019.

Chou, A., Morganti, J.M., and Rosi, S. (2016). Frontal Lobe Contusion in Mice Chronically Impairs Prefrontal-Dependent Behavior. PLoS One 11, e0151418. 10.1371/journal.pone.0151418.

Cobos, I., Calcagnotto, M.E., Vilaythong, A.J., Thwin, M.T., Noebels, J.L., Baraban, S.C., and Rubenstein, J.L. (2005). Mice lacking Dlx1 show subtype-specific loss of interneurons, reduced inhibition and epilepsy. Nat Neurosci 8, 1059–1068. 10.1038/nn1499.

Cossart, R., Dinocourt, C., Hirsch, J.C., Merchan-Perez, A., De Felipe, J., Ben-Ari, Y., Esclapez, M., and Bernard, C. (2001). Dendritic but not somatic GABAergic inhibition is decreased in experimental epilepsy. Nat Neurosci 4, 52–62. 10.1038/82900.

Cottam, J.C., Smith, S.L., and Hausser, M. (2013). Target-specific effects of somatostatin-expressing interneurons on neocortical visual processing. J Neurosci 33, 19567–19578. 10.1523/JNEUROSCI.2624-13.2013.

Ding, M.C., Wang, Q., Lo, E.H., and Stanley, G.B. (2011). Cortical excitation and inhibition following focal traumatic brain injury. J Neurosci 31, 14085–14094. 10.1523/JNEUROSCI.3572-11.2011.

Engberg, A.W., and Teasdale, T.W. (2004). Psychosocial outcome following traumatic brain injury in adults: a long-term population-based follow-up. Brain Inj 18, 533–545. 10.1080/02699050310001645829.

Folweiler, K.A., Xiong, G., Best, K.M., Metheny, H.E., Nah, G., and Cohen, A.S. (2020). Traumatic Brain Injury Diminishes Feedforward Activation of Parvalbumin-Expressing Interneurons in the Dentate Gyrus. eNeuro 7. 10.1523/ENEURO.0195-19.2020.

Fujiwara, E., Schwartz, M.L., Gao, F., Black, S.E., and Levine, B. (2008). Ventral frontal cortex functions and quantified MRI in traumatic brain injury. Neuropsychologia 46, 461–474. 10.1016/j.neuropsychologia.2007.08.027.

Goodwill, H.L., Manzano-Nieves, G., LaChance, P., Teramoto, S., Lin, S., Lopez, C., Stevenson, R.J., Theyel, B.B., Moore, C.I., Connors, B.W., and Bath, K.G. (2018). Early Life Stress Drives Sex-Selective Impairment in Reversal Learning by Affecting Parvalbumin Interneurons in Orbitofrontal Cortex of Mice. Cell Rep 25, 2299–2307 e2294. 10.1016/j.celrep.2018.11.010.

Griffin, A., Hamling, K.R., Hong, S., Anvar, M., Lee, L.P., and Baraban, S.C. (2018). Preclinical Animal Models for Dravet Syndrome: Seizure Phenotypes, Comorbidities and Drug Screening. Front Pharmacol 9, 573. 10.3389/fphar.2018.00573.

Guerriero, R.M., Giza, C.C., and Rotenberg, A. (2015). Glutamate and GABA imbalance following traumatic brain injury. Curr Neurol Neurosci Rep 15, 27. 10.1007/s11910-015-0545-1.

Ito-Ishida, A., Ure, K., Chen, H., Swann, J.W., and Zoghbi, H.Y. (2015). Loss of MeCP2 in Parvalbumin-and Somatostatin-Expressing Neurons in Mice Leads to Distinct Rett Syndrome-like Phenotypes. Neuron 88, 651–658. 10.1016/j.neuron.2015.10.029.

Mao, R., Page, D.T., Merzlyak, I., Kim, C., Tecott, L.H., Janak, P.H., Rubenstein, J.L., and Sur, M. (2009). Reduced conditioned fear response in mice that lack Dlx1 and show subtype-specific loss of interneurons. J Neurodev Disord 1, 224–236. 10.1007/s11689-009-9025-8.

Murayama, M., Perez-Garci, E., Nevian, T., Bock, T., Senn, W., and Larkum, M.E. (2009). Dendritic encoding of sensory stimuli controlled by deep cortical interneurons. Nature 457, 1137–1141. 10.1038/nature07663.

Naka, A., and Adesnik, H. (2016). Inhibitory Circuits in Cortical Layer 5. Front Neural Circuits 10, 35. 10.3389/fncir.2016.00035.

Naka, A., Veit, J., Shababo, B., Chance, R.K., Risso, D., Stafford, D., Snyder, B., Egladyous, A., Chu, D., Sridharan, S., et al. (2019). Complementary networks of cortical somatostatin interneurons enforce layer specific control. Elife 8. 10.7554/eLife.43696.

Nichols, J., Bjorklund, G.R., Newbern, J., and Anderson, T. (2018). Parvalbumin fast-spiking interneurons are selectively altered by paediatric traumatic brain injury. J Physiol 596, 1277–1293. 10.1113/JP275393.

Nigro, M.J., Hashikawa-Yamasaki, Y., and Rudy, B. (2018). Diversity and Connectivity of Layer 5 Somatostatin-Expressing Interneurons in the Mouse Barrel Cortex. J Neurosci 38, 1622–1633. 10.1523/JNEUROSCI.2415-17.2017.

Ponsford, J., Draper, K., and Schonberger, M. (2008). Functional outcome 10 years after traumatic brain injury: its relationship with demographic, injury severity, and cognitive and emotional status. J Int Neuropsychol Soc 14, 233–242. 10.1017/S1355617708080272.

Potter, G.B., Petryniak, M.A., Shevchenko, E., McKinsey, G.L., Ekker, M., and Rubenstein, J.L. (2009). Generation of Cre-transgenic mice using Dlx1/Dlx2 enhancers and their characterization in GABAergic interneurons. Mol Cell Neurosci 40, 167–186. 10.1016/j.mcn.2008.10.003.

Pouille, F., Marin-Burgin, A., Adesnik, H., Atallah, B.V., and Scanziani, M. (2009). Input normalization by global feedforward inhibition expands cortical dynamic range. Nat Neurosci 12, 1577–1585. 10.1038/nn.2441.

Pouille, F., and Scanziani, M. (2001). Enforcement of temporal fidelity in pyramidal cells by somatic feed-forward inhibition. Science 293, 1159–1163. 10.1126/science.1060342.

Raible, D.J., Frey, L.C., Cruz Del Angel, Y., Russek, S.J., and Brooks-Kayal, A.R. (2012). GABA(A) receptor regulation after experimental traumatic brain injury. J Neurotrauma 29, 2548–2554. 10.1089/neu.2012.2483.

Riedemann, T. (2019). Diversity and Function of Somatostatin-Expressing Interneurons in the Cerebral Cortex. Int J Mol Sci 20. 10.3390/ijms20122952.

Rudy, B., Fishell, G., Lee, S., and Hjerling-Leffler, J. (2011). Three groups of interneurons account for nearly 100% of neocortical GABAergic neurons. Dev Neurobiol 71, 45–61. 10.1002/dneu.20853.

Silberberg, G., and Markram, H. (2007). Disynaptic inhibition between neocortical pyramidal cells mediated by Martinotti cells. Neuron 53, 735–746. 10.1016/j.neuron.2007.02.012.

Spikman, J.M., Timmerman, M.E., Milders, M.V., Veenstra, W.S., and van der Naalt, J. (2012). Social cognition impairments in relation to general cognitive deficits, injury severity, and prefrontal lesions in traumatic brain injury patients. J Neurotrauma 29, 101–111. 10.1089/neu.2011.2084.

Struchen, M.A., Clark, A.N., Sander, A.M., Mills, M.R., Evans, G., and Kurtz, D. (2008). Relation of executive functioning and social communication measures to functional outcomes following traumatic brain injury. NeuroRehabilitation 23, 185–198.

Stuss, D.T. (2011). Traumatic brain injury: relation to executive dysfunction and the frontal lobes. Curr Opin Neurol 24, 584–589. 10.1097/WCO.0b013e32834c7eb9.

Tai, C., Abe, Y., Westenbroek, R.E., Scheuer, T., and Catterall, W.A. (2014). Impaired excitability of somatostatin- and parvalbumin-expressing cortical interneurons in a mouse model of Dravet syndrome. Proc Natl Acad Sci U S A 111, E3139–3148. 10.1073/pnas.1411131111.

Um, J.W. (2017). Roles of Glial Cells in Sculpting Inhibitory Synapses and Neural Circuits. Front Mol Neurosci 10, 381. 10.3389/fnmol.2017.00381.

Vascak, M., Jin, X., Jacobs, K.M., and Povlishock, J.T. (2018). Mild Traumatic Brain Injury Induces Structural and Functional Disconnection of Local Neocortical Inhibitory Networks via Parvalbumin Interneuron Diffuse Axonal Injury. Cereb Cortex 28, 1625–1644. 10.1093/cercor/bhx058.

Vascak, M., Sun, J., Baer, M., Jacobs, K.M., and Povlishock, J.T. (2017). Mild Traumatic Brain Injury Evokes Pyramidal Neuron Axon Initial Segment Plasticity and Diffuse Presynaptic Inhibitory Terminal Loss. Front Cell Neurosci 11, 157. 10.3389/fncel.2017.00157.

Vilkki, J., Ahola, K., Holst, P., Ohman, J., Servo, A., and Heiskanen, O. (1994). Prediction of psychosocial recovery after head injury with cognitive tests and neurobehavioral ratings. J Clin Exp Neuropsychol 16, 325–338. 10.1080/01688639408402643.

Wallesch, C.W., Curio, N., Galazky, I., Jost, S., and Synowitz, H. (2001). The neuropsychology of blunt head injury in the early postacute stage: effects of focal lesions and diffuse axonal injury. J Neurotrauma 18, 11–20. 10.1089/089771501750055730.

Wang, X.J., Tegner, J., Constantinidis, C., and Goldman-Rakic, P.S. (2004a). Division of labor among distinct subtypes of inhibitory neurons in a cortical microcircuit of working memory. Proc Natl Acad Sci U S A 101, 1368–1373. 10.1073/pnas.0305337101.

Wang, Y., Toledo-Rodriguez, M., Gupta, A., Wu, C., Silberberg, G., Luo, J., and Markram, H. (2004b). Anatomical, physiological and molecular properties of Martinotti cells in the somatosensory cortex of the juvenile rat. J Physiol 561, 65–90. 10.1113/jphysiol.2004.073353.

Willems, J.G.P., Wadman, W.J., and Cappaert, N.L.M. (2018). Parvalbumin interneuron mediated feedforward inhibition controls signal output in the deep layers of the perirhinal-entorhinal cortex. Hippocampus 28, 281–296. 10.1002/hipo.22830.

Wilson, L., Stewart, W., Dams-O’Connor, K., Diaz-Arrastia, R., Horton, L., Menon, D.K., and Polinder, S. (2017). The chronic and evolving neurological consequences of traumatic brain injury. Lancet Neurol 16, 813–825. 10.1016/S1474-4422(17)30279-X.

Xu, H., Jeong, H.Y., Tremblay, R., and Rudy, B. (2013). Neocortical somatostatin-expressing GABAergic interneurons disinhibit the thalamorecipient layer 4. Neuron 77, 155–167. 10.1016/j.neuron.2012.11.004.

Yang, Z., Wu, G., Liu, M., Sun, X., Xu, Q., Zhang, C., and Lei, H. (2021). Dysfunction of Orbitofrontal GABAergic Interneurons Leads to Impaired Reversal Learning in a Mouse Model of Obsessive-Compulsive Disorder. Curr Biol 31, 381–393 e384. 10.1016/j.cub.2020.10.045.

Zhu, Y., Qiao, W., Liu, K., Zhong, H., and Yao, H. (2015). Control of response reliability by parvalbumin-expressing interneurons in visual cortex. Nat Commun 6, 6802. 10.1038/ncomms7802.

