## Supplemental Figures and Tables for "Selective Inhibitory Circuit Dysfunction after Chronic Frontal Lobe Contusion"

**Supplemental Table 1. Intrinsic Properties of Layer V Neurons.** Values are mean (SEM) or *median (95% CI). Statistical test is indicated with associated p-value.

| Variables | non-FS Sham | non-FS TBI | *p-value* | FS Sham | FS TBI | *p-value* | PYR Sham | PYR TBI | *p-value* |
|---|---|---|---|---|---|---|---|---|---|
| **Action potential properties** | | | | | | | | | |
| half width (msec) | 0.56 (0.03) | 0.65 (0.02) | ***0.0053*** (unpaired t-test) | 0.29 (0.01) | 0.27 (0.01) | *0.2574* (unpaired t-test) | 0.86 (0.07) | 0.78 (0.04) | *0.2938* (unpaired t-test) |
| amplitude (mV) | 52.07 (2.90) | 55.00 (2.68) | *0.4808* (unpaired t-test) | 53.75 (1.80) | 54.75 (2.01) | *0.7432* (unpaired t-test) | 74.12 (2.26) | 75.79 (1.32) | *0.5154* (unpaired t-test) |
| threshold (mV) | -36.09 (1.18) | -36.18 (1.02) | *0.9529* (unpaired t-test) | -38.61(1.22) | -41.37 (1.52) | *0.1614* (unpaired t-test) | -39.56 (1.38) | -37.53 (1.08) | *0.2491* (unpaired t-test) |
| rising slope (mV/s) | *191.8 (165.4 - 212.7) | *177.3 (162.4 - 236.5) | *0.7366* (Mann-Whitney) | 302.5 (17.21) | 320.5 (12.87) | *0.4253* (unpaired t-test) | 301.8 (19.99) | 353.8 (13.73) | ***0.0368*** (unpaired t-test) |
| falling slope (mV/s) | -116.6 (9.38) | -94.68 (4.12) | *0.0530* (Welch's t-test) | -252.2 (19.13) | -286.2 (16.77) | *0.2010* (unpaired t-test) | -99.33 (11.03) | -92.38 (6.77) | *0.5855* (unpaired t-test) |
| spike AHP (mV) | 15.11 (1.41) | 13.42 (0.64) | *0.2323* (unpaired t-test) | 22.61 (0.62) | 21.85 (1.01) | *0.5087* (unpaired t-test) | 11.59 (0.69) | 15.59 (0.89) | ***0.0015*** (unpaired t-test) |
| adaptation index | 2.35 (0.16) | 3.25 (0.33) | ***0.0226*** (Welch's t-test) | 1.40 (0.04) | 1.292 (0.04) | *0.0994* (unpaired t-test) | *2.59 (2.51 - 3.94) | *2.43 (2.12 - 3.48) | *0.2777* (Mann-Whitney) |
| rheobase (mV) | 69.76 (20.73) | 58.75 (16.76) | *0.6849* (unpaired t-test) | 194.20 (31.67) | 194 (28.01) | *0.9971* (unpaired t-test) | 86.31 (14.48) | 86.1 (12.32) | *0.9915* (unpaired t-test) |
| **Passive properties** | | | | | | | | | |
| resting potential (mV) | -61.3 (2.92) | -59.19 (1.77) | *0.5166* (unpaired t-test) | -65.12 (1.40) | -66.71 (1.85) | *0.4899* (unpaired t-test) | -64.53 (1.85) | -66.06 (1.48) | *0.5200* (unpaired t-test) |
| membrane resistance (MΩ) | 271.3 (37.06) | 279.2 (26.77) | *0.8618* (unpaired t-test) | 140.6 (16.22) | 124.8 (13.34) | *0.4702* (unpaired t-test) | 133.3 (16.37) | 130.8 (10.60) | *0.8961* (unpaired t-test) |
| tau (msec) | 15.67 (2.18) | 19.08 (2.70) | *0.3826* (unpaired t-test) | *7.55 (6.10 - 9.53) | *6.9 (5.79 - 8.54) | *0.4041* (Mann-Whitney) | 17.7 (1.77) | 20.97 (1.63) | *0.1836* (unpaired t-test) |

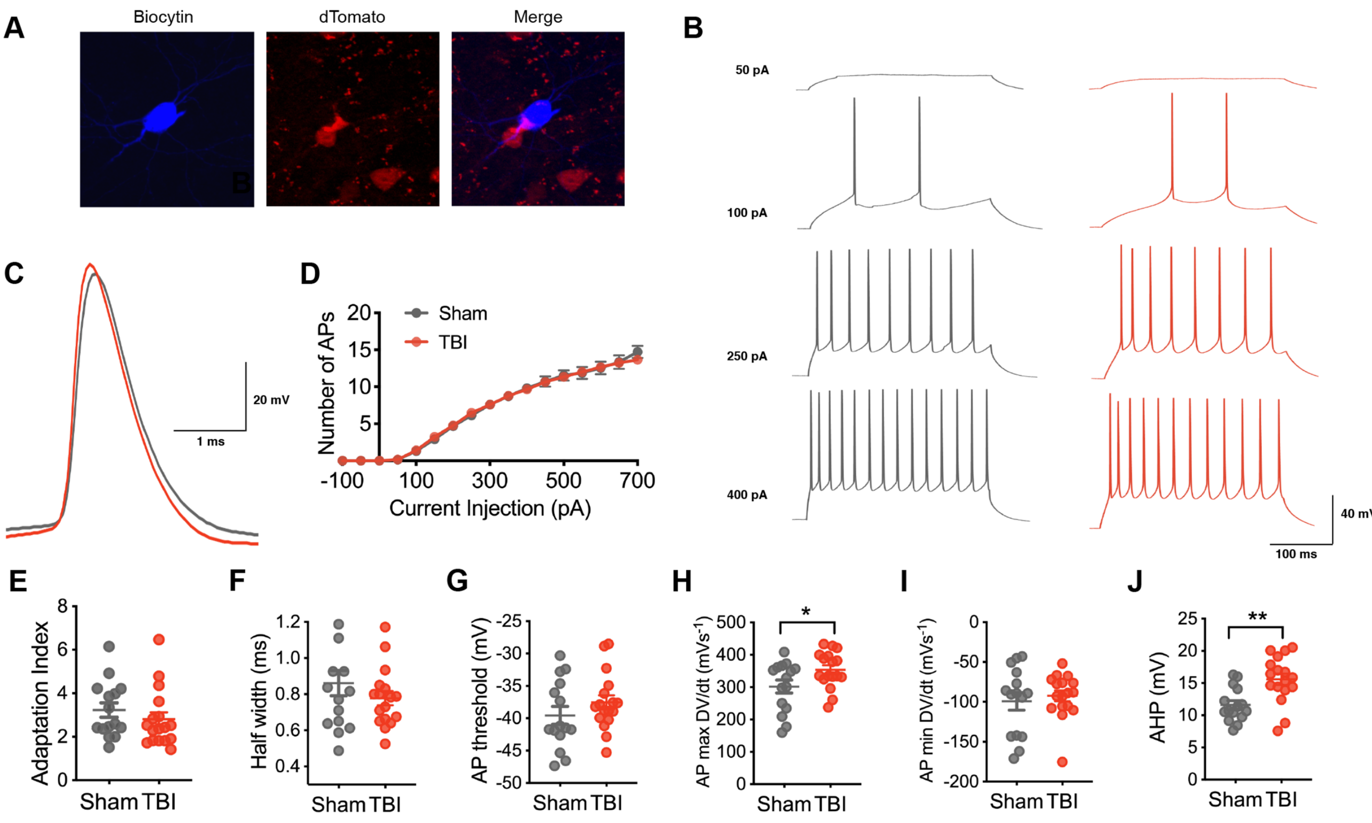


**Supplemental Figure 1. Intrinsic excitability in layer V pyramidal neurons after chronic TBI.** A) A pyramidal neuron in layer V of the orbitofrontal cortex that was filled with biocytin (blue) and later confirmed to not express tdTomato (red). B) Representative current-clamp responses to depolarizing current steps in sham (grey) and TBI (red) mice. C) Average action potential shape. D) The number of action potentials plotted as a function of current injection ($p = 0.9608$ for TBI effect, repeated measures 2-way ANOVA). E) The adaptation index from current clamp responses measured at 400 pA above spiking threshold ( $p = 0.2777$, Mann-Whitney test). F-J) The action potential (AP) half-width, AP threshold, AP maximum rising slope, AP minimum falling slope, and afterhyperpolarization (AHP) calculated from current clamp responses 100 pA above spiking threshold ($p = 0.2938$, unpaired t-test; $p = 0.2491$, unpaired t-test; * $p = 0.0368$, unpaired t-test; $p = 0.5855$, unpaired t-test; ** $p = 0.0015$, unpaired t-test; respectively).

Each neuron is represented with a symbol; solid lines indicate the mean ± SEM. (n=15 sham and 17 TBI layer V pyramidal neurons from 6 (sham) and 7 (TBI) animals/group).

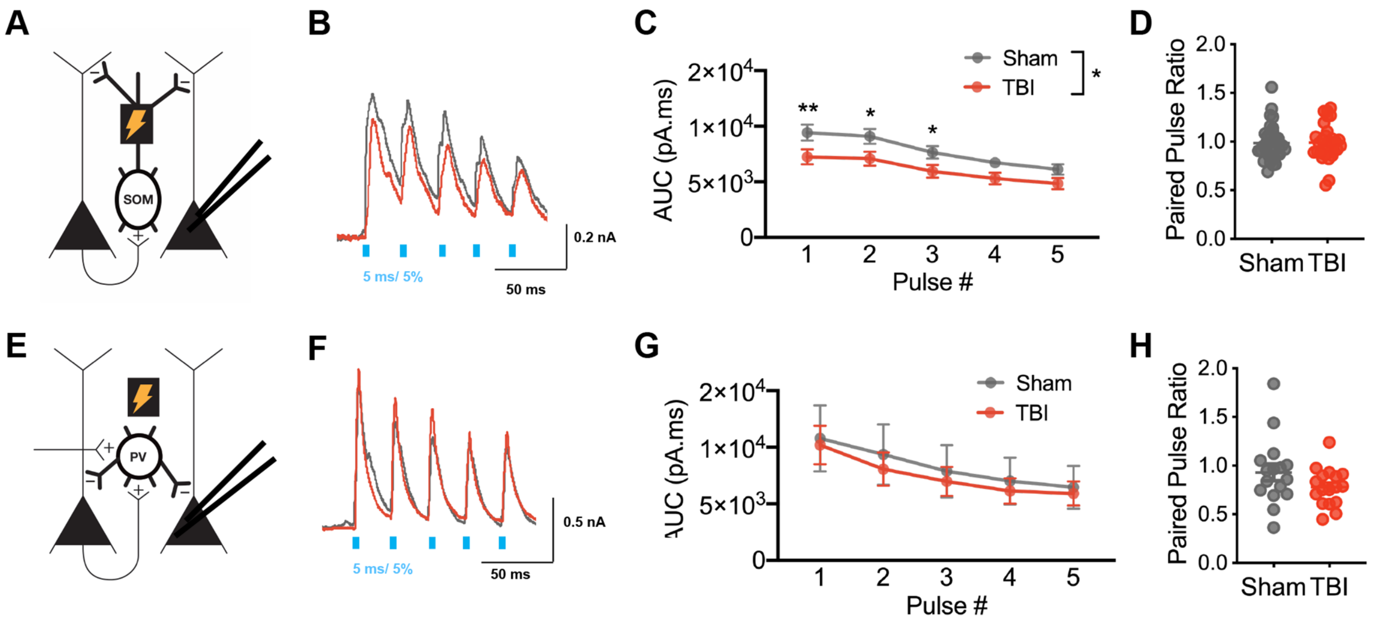


**Supplemental Figure 2. Selective reduced SOM+-mediated inhibitory synaptic input in layer V pyramidal neurons after TBI with 40 Hz repetitive stimulation.** A) Schematic of experimental design: voltage clamp recordings were obtained from layer V pyramidal neurons while activating ChR2-expressing SOM+ interneurons. B) Example oIPSCs in sham (grey) and TBI (red) conditions elicited with 40 Hz stimulation. C) Total charge (AUC) of oIPSCs elicited across pulse number (* p = 0.0374 for TBI effect, repeated measures two-way ANOVA; ** p = 0.0093, * $p < 0.05$ post hoc tests controlling for the false discovery rate). D) The paired pulse ratio determined as the ratio of pulse 2/pulse 1 (p = 0.9064, unpaired t-test). E) Schematic of experimental design: voltage clamp recordings were obtained from layer V pyramidal neurons while activating ChR2- expressing PV+ interneurons. F) Example oIPSCs in sham (grey) and TBI (red) conditions elicited with 40 Hz stimulation. G) Total charge (AUC) of oIPSC elicited across pulse number (p = 0.7540 for TBI effect, repeated measures two-way ANOVA). H) The paired pulse ratio determined as the ratio of pulse 2/pulse 1 (p = 0.1364; unpaired t-test with Welch's correction).

Circles represent the mean, solid lines indicate the SEM in C, G; Each neuron/slice is represented with a symbol and solid lines indicate the mean ± SEM in D, H, I. (n=34 sham and 30 TBI neurons from 7 (sham) and 6 (TBI) animals/group for oIPSC data from SOM+ stimulation; n= 16 sham and 17 TBI neurons from 4 (sham) and 4 (TBI) animals/group for oIPSC data from PV+ stimulation).

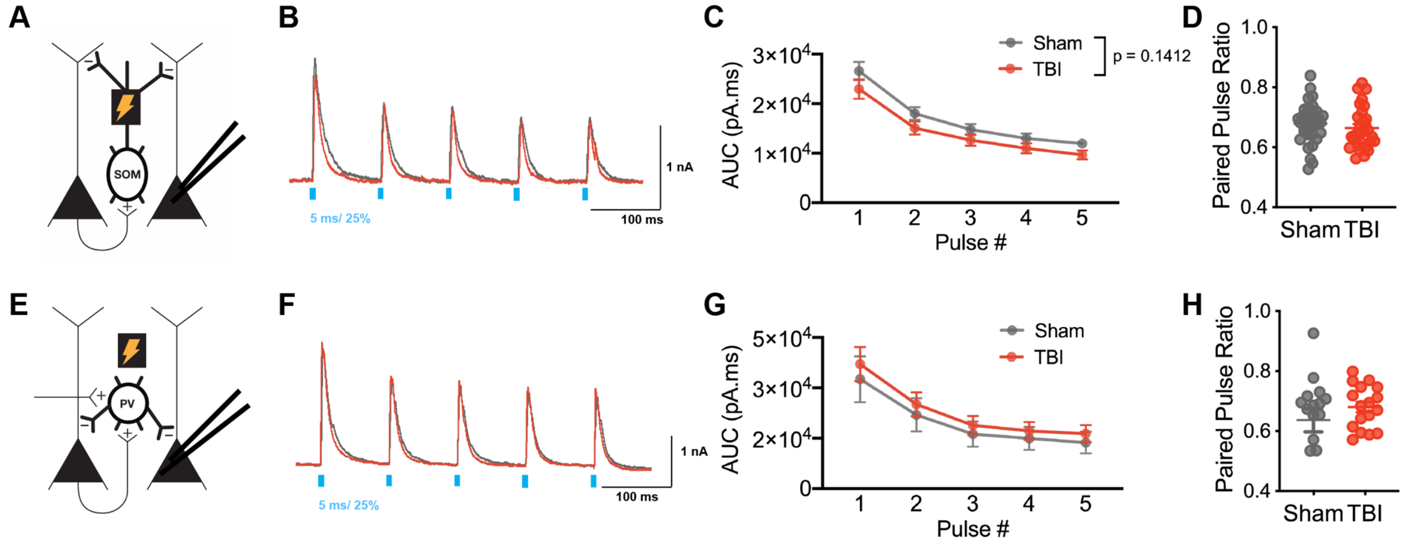


**Supplemental Figure 3. SOM+- and PV+-mediated inhibitory synaptic input in layer V pyramidal neurons with 10 Hz repetitive stimulation at 25% light intensity.** A) Schematic of experimental design: voltage clamp recordings were obtained from layer V pyramidal neurons while activating ChR2-expressing SOM+ interneurons. B) Example oIPSCs in sham (grey) and TBI (red) conditions elicited with 10 Hz stimulation at 25% light intensity. C) Total charge (AUC) of oIPSCs elicited across pulse number ($p = 0.1412$ for TBI effect, repeated measures two-way ANOVA. D) The paired pulse ratio determined as the ratio of pulse 2/pulse 1 ($p = 0.4160$, unpaired t-test). E) Schematic of experimental design: voltage clamp recordings were obtained from layer V pyramidal neurons while activating ChR2- expressing PV+ interneurons. F) Example oIPSCs in sham (grey) and TBI (red) conditions elicited with 10 Hz stimulation at 25% light intensity. G) Total charge (AUC) of oIPSC elicited across pulse number ($p = 0.5855$ for TBI effect, repeated measures two-way ANOVA). H) The paired pulse ratio determined as the ratio of pulse 2/pulse 1 ($p = 0.3199$; unpaired t-test with Welch's correction).

Circles represent the mean, solid lines indicate the SEM in C, G; Each neuron/slice is represented with a symbol and solid lines indicate the mean ± SEM in D, H, I. (n=34 sham and 30 TBI neurons from 7 (sham) and 6 (TBI) animals/group for oIPSC data from SOM+ stimulation; n= 16 sham and 17 TBI neurons from 4 (sham) and 4 (TBI) animals/group for oIPSC data from PV+ stimulation).
