## Supplemental Key Resource Table for "Selective Inhibitory Circuit Dysfunction after Chronic Frontal Lobe Contusion"

**KEY RESOURCES TABLE**

| REAGENT or RESOURCE | SOURCE | IDENTIFIER |
| --- | --- | --- |
| **Antibodies** | | |
| Rabbit anti-somatostatin | Abcam | Cat.#: AB64053; RRID:AB_1143012 |
| Mouse anti-parvalbumin | Millipore | Cat.#:MAB1572; RRID:AB_2174013 |
| Anti-Rabbit Alexafluor488 | Molecular Probes (Invitrogen) | Cat.#: A21206; RRID:AB_2535792 |
| Anti-Mouse Alexafluor647 | Abcam | Cat.#: AB181292 |
| Rabbit anti-GFP | EXBIO | Cat.#: 11-476; RRID:AB_10735170 |
| **Chemicals, peptides, and recombinant proteins** |  |  |
| streptavidin-conjugated Pacific Blue | ThermoFisher Scientific | Cat.#: S11222 |
| **Experimental models: Organisms/strains** |  |  |
| PV-Cre | The Jackson Laboratory | Cat.#: 017320; RRID:IMSR_JAX:017320 |
| SOM-Cre | The Jackson Laboratory | Cat.#: 013044; RRID:IMSR_JAX:013044 |
| DlxI12b-Cre | Dr. John Rubenstein | NA |
| Ai14(B6.Cg-Gt(ROSA)26Sor^tm14(CAG-tdTomato)Hze^/J) | The Jackson Laboratory | Cat.#: 007914 RRID:IMSR_JAX:007914 |
| Ai32(B6.Cg-Gt(ROSA)26Sor^tm32(CAG-COP4*H134R/EYFP)Hze^/J) | The Jackson Laboratory | Cat.#: 024109 RRID:IMSR_JAX:024109 |
| **Software and algorithms** |  |  |
| ClampFit | Molecular Devices | RRID:SCR_011323 |
| ImageJ | National Center for Microscopy and Imaging Research | RRID:SCR_001935 |
| Zen Software | Zeiss | RRID:SCR_018163 |
| Graph Pad Prism | Graph Pad | RRID:SCR_002798 |
